# The essential role played by B cells in supporting protective immunity against *Trichuris muris* infection is dependent on host genetic background and is independent of antibody

**DOI:** 10.1101/550434

**Authors:** Rinal Sahputra, Dominik Ruckerl, Kevin Couper, Werner Muller, Kathryn J Else

**Affiliations:** Lydia Becker Institute for Immunology, The University of Manchester

## Abstract

This study investigates the role of B cells in immunity to *Trichuris muris* (*T. muris*) infection in two genetically distinct strains of mouse, using anti-CD20 monoclonal antibody (mAb) (Genentech-clone 5D2) to deplete B cells. Data is presented for the mouse strains: C57BL/6 and BALB/c, which mount mixed Th1/Th2, and highly polarised Th2 immune responses to *T. muris*, respectively. C57BL/6 mice receiving anti-CD20 treatment prior to and during, or anti-CD20 treatment that commenced two weeks post infection (p.i.), were susceptible to *T. muris*. Parasite-specific IgG1 antibodies were absent and Th2 type cytokines produced by mesenteric lymph nodes cells from mice receiving α-CD20 mAb treatment were significantly lower than produced by cells from isotype control treated mice. T follicular helper cells were also significantly reduced. Importantly, and in complete contrast, BALB/c mice were still able to expel *T.muris* in the absence of B cells, revealing that the essential role played by B cells in protective immunity was dependent on genetic background. To explore whether the important role played by the B cell in the protective immune response of C57BL/6 mice was in enabling strong Th2 responses in the presence of IFN-γ, IFN-γ was blocked using anti-IFN-γ mAb post B cell depletion. Depleting IFN-γ, in the absence of B cells restored worm expulsion in the absence of parasite-specific IgG1/IgG2c and partially rescued the *T. muris* specific IL-13 response. Thus, our data suggest an important, antibody independent role for B cells in supporting Th2 type immune responses in mixed IFN-γ-rich Th1/Th2 immune response settings.

**Author summary:** How B cells contribute to protective immunity against parasitic nematodes remains unclear, with their importance as accessory cells under-explored. This study reveals that, on some genetic backgrounds, B cells are important for the expulsion of *T. muris* by acting as accessory cells, supporting Th2 immune responses.

## Introduction

Infecting over two billion people around the world, mostly in resource-limited countries, the ability of parasitic helminths to maintain long standing chronic infections makes them a major health care issue (1). *Trichuris trichiura* (*T. trichiura*) is one of the most common gastrointestinal nematodes, infecting approximately 465 million people worldwide (2), primarily children. In infected children, trichuriasis is strongly associated to malnutrition, growth stunting, and intellectual retardation; whereas in pregnant adults it is related to anaemia and low birth weight babies (3).

For decades, *T. muris* in the mouse has provided a useful and relevant model system with which to explore immunity to *T. trichiura* in man. Infection of mice with intestinal nematode parasite *T. muris* drives polarized T helper cell (Th) responses which associate with resistance (Th2) or susceptibility (Th1). However, the key cellular contributions which support Th2 cell polarization during *T. muris* infection are still not well understood. One of the cells thought to be important is the B cell. It is well established that B cell function is not only related to antibody production, but B cells can also act as antigen presenting cells (APCs) due to the expression of MHC class II molecules and several co-stimulators, including CD40, CD80, and CD86 on its surface (4–6), and as accessory cells, acting as a cellular source of multiple cytokines (7).

B cells are well placed to act as accessory cells as they are able to influence CD4 T cells polarisation. For example, previous studies have shown that CD4 T cells produce significantly more IL-4 when stimulated with antigen presented by B cells compared to macrophages (8). In addition, B cells have been shown to produce either Th1 or Th2 type cytokines *in vivo* depending on the type of parasite. Thus, for the Th1 driving parasite *Toxoplasma gondii*, B cells produce IFN-γ and IL-12p40 (9); whilst in the context of the Th2 polarising parasite *Heligmosomoides polygyrus* (*H. polygyrus*), B cells produce IL-4 and IL-2 (10, 11). Recent studies reported that IL-4 producing B cells during early infection of *Schistosoma mansoni* (*S. mansoni*) are critical for Th2 polarisation, protecting the host against *S. mansoni*-induced pathology (12). B cells can also influence T cell polarisation towards Th2 response by providing co-stimulatory molecules. Linton *et al*., (13) showed that co-stimulation via OX40L expressed by B cells is essential for T cell polarisation towards Th2 cells. Likewise, both CD80 and CD86 co-stimulatory molecules expressed by B cells, as well as other APC, are required for the Th2 response during *H. polygyrus* infection (14).

Recently, it was suggested that B cells, together with T cells and DCs are required not only to optimize the development of Th2 type responses, but also to maintain T follicular helper (T_FH_) and memory Th2 cells development following *H. polygyrus* infection (15). Thus, Leon *et al.* (15) showed that the deletion of CXCR5 on either DC or CD4 T cells, during *H. polygyrus*-infection of C57BL/6 mice, impaired the development of T_FH_ and Th2 cells by impairing the migration of CXCR5+ cells towards the B cell follicle in response to CXCL13. Interestingly, B cell depletion also impaired Th2 responses (15).

Previous studies suggested that B cells and antibody are not important in mediating resistance following a primary *T. muris* infection (16–18). Since then, many different studies have been performed to show the importance of CD4 T cells in mediating resistance against *T. muris* (19–21). By contrast, the role of B cells in immunity to *T. muris* remains largely unexplored. *T. muris*-infected µMT mice on a C57BL6 background develop Th-1 type responses, resulting in the susceptibility to *T. muris* infection (22). These susceptible *T. muris*-infected µMT C57BL6 mice produced high IFN-γ, without any Th2 cytokines production. Furthermore, when these mice were treated with B cells from naive C57BL6 or with anti-IL-12 antibody, resistance to infection was restored (22). This suggests that B cells are important in either inhibiting Th1 development or supporting Th2 type immune responses. However, given the importance of B cells in the development of lymph nodes and tissue organisation (23, 24), data from µMT mice is difficult to interpret.

This study therefore investigates the role of B cells and antibodies in immunity to *T. muris* infection using anti-CD20 monoclonal antibody (mAb) to deplete B cells from mice of two distinct genetic backgrounds, C57BL/6 and BALB/c. The benefit of using anti-CD20 mAb is that it allows depletion of CD19+ cells either prior to or post infection and avoids the complicating consequences of B cell deficiency during embryonic development. We demonstrate that B cells are important in the protective immune response to *T. muris*; that the role played by the B cell is antibody-independent; and that the importance of the B cell varies with genetic background of the host. Thus the B cell plays an essential role in supporting Th2 type immune responses only in mixed Th1/Th2 IFN-γ rich settings, as seen in C57BL/6 mice, and is redundant in the highly Th2 polarised, IFN−γ-deficient environment of the BALB/c mouse post *T. muris* infection.

## Results

### 2.1 Depletion of B cells throughout infection requires two injections of anti-CD20 mAb

As a transmembrane calcium channel, CD20 is important in B cell activation, proliferation, and differentiation (25). The CD20 molecule is normally expressed on the surface of B cells during the late pre-B cell. Therefore, the injection of anti-CD20 mAb will deplete all B cells, except early pre-B cells and plasma cells (26). The vast majority of IgM^+^ or B220^+^ cells in the spleen, lymph nodes, peritoneal cavity, and blood express CD20, whilst in bone marrow the expression of CD20 increases with B cell maturation (27). Previous studies showed that anti-CD20 effectively depleted B cells either via complement-mediated lysis of target cells (28) or by inducing FcγR-mediated clearance (Ab-dependent cell mediated cytotoxicity) (29). Furthermore, a single dose of anti-CD20 treatment prior to lymphocytic choriomeningitis virus infection depleted B cell populations for up to 45 days (30). However, the current study showed that a single anti-CD20 injection failed to ablate B cells for the full duration of *T. muris* infection, with a clear CD19+ population re-emerging in the MLNs, spleen, and blood by day 35 p.i. (Fig. 1A&B). Therefore, a second anti-CD20 injection was administered at day 10 p.i. in order to maintain B cell depletion in the blood and secondary lymphoid organs throughout infection (Fig. 1A&B). CD19^+^ cells were still present in the bone marrow after anti-CD20 treatment (Fig. 1B), however these CD19^+^ cells were pro-B cells (Fig. 1C) as has been shown previously (26).

**Fig. 1.**
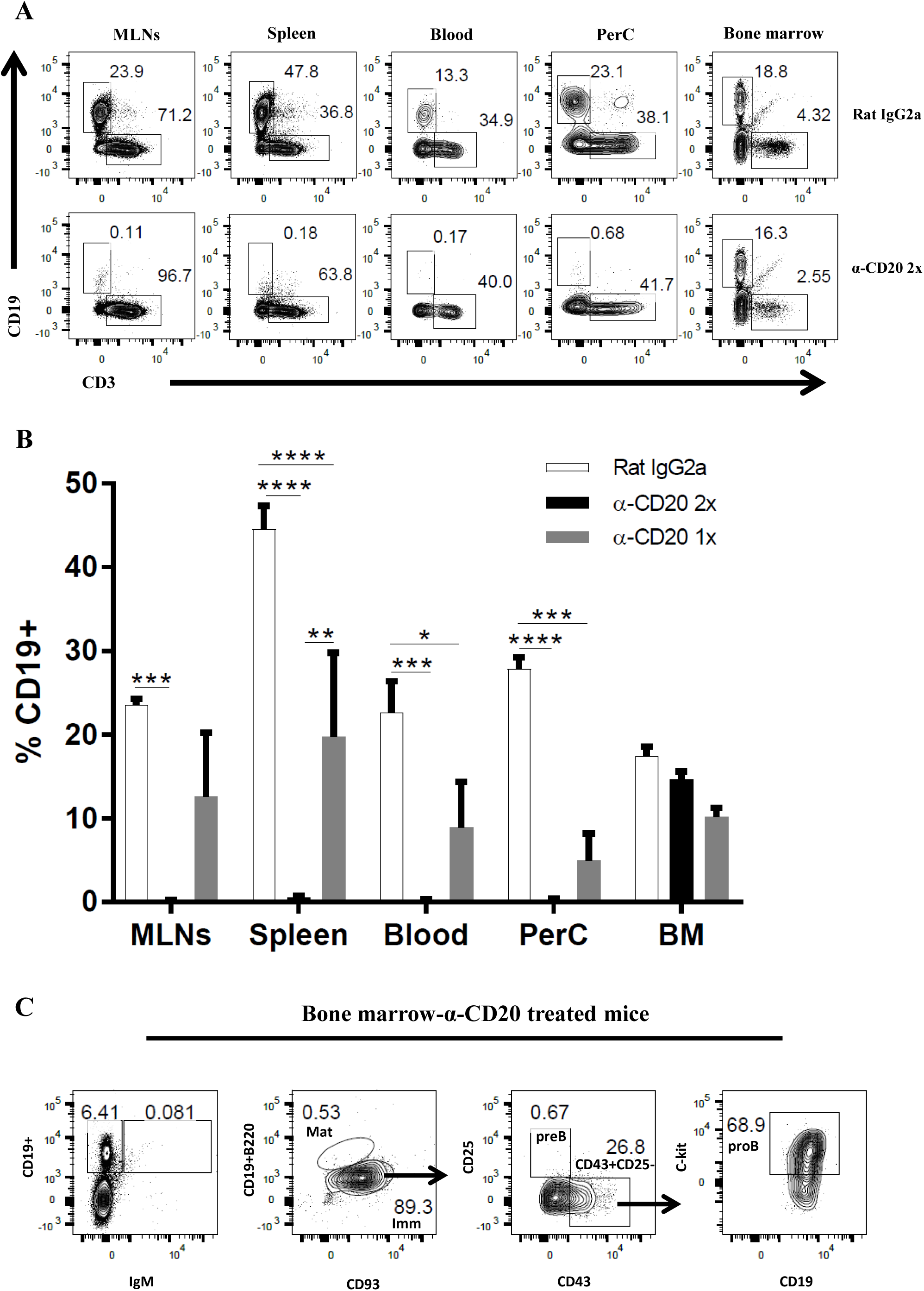
Two injections of anti CD20-mab are required to ensure B cells are depleted occurred throughout infection. C57BL/6 mice were treated with anti CD20 mAb or Rat IgG2a isotype control at 100 μg in 200 μl PBS by i.v. injection via tail vein. Mice were infected with approximately 150 *T. muris* eggs at day 7 post injection. Some mice were only injected once with anti CD20/ rat IgG2a isotype control, whilst some mice were re-injected i.v. with anti CD20 mAb or Rat IgG2a at 100 μg in 200 μl PBS on day 10 p.i. Mice were autopsied at day 35 p.i. (A&B) CD19+ cells were assessed in MLNs, spleen, blood, peritoneal cavity (PerC), and bone marrow (BM) using flow cytometry. (C) Representative flow cytometric analysis of CD19+ cells in BM of anti-CD20 treated mice. Both immature and mature B cells are IgM+, whilst pro/pre-B cells are IgM-. Pro-B cells are defined as CD19+B220^low^IgM-CD93+CD43+CD25-ckit+. Data shown is mean ± SEM, representative of 2 independent experiments for Rat IgG2a and α-CD20 2x, from 1 experiment for α-CD20 1x, n=5, males. *p<0.05, **p<0.01, ***p<0.001, ****p<0.0001, Anova.

### 2.2 C57BL/6 mice treated with anti-CD20 mAb fail to expel *T. muris* by d35 p.i., correlates with an absence of class switched antibodies and a significant decrease in Th2 cytokines

To investigate whether B cells are important in resistance against *T. muris* during primary infection of C57BL/6 mice, mice were injected with anti-CD20 mAb 7 days prior to infection and 10 days post infection. Chronic infection, characterised by persisting adult stage parasites from day 32 p. i. defines susceptibility (22). Thus, autopsies were performed beyond this time point. The experimental design is shown in Fig. 2A. Previous studies have shown that C57BL/6 mice take up to 35 days to completely expel the parasite (31). As shown in Fig. 2B, C57BL/6 depleted of B cells were significantly more susceptible to infection than control treated mice.

**Fig. 2.**
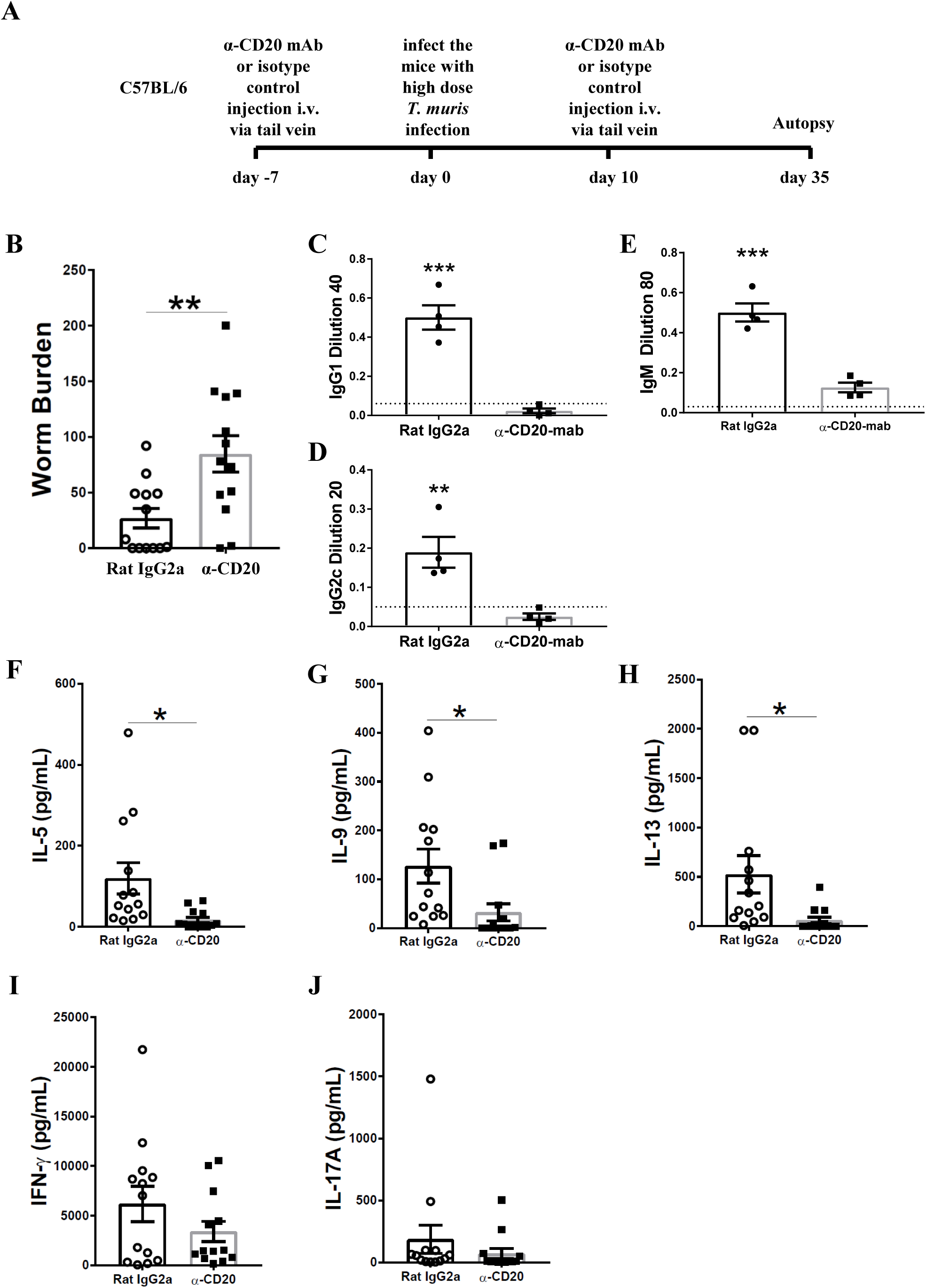
B cell depleted C57BL6 mice were susceptible to *T. muris*, correlating with an absence of class switched antibodies and a significant decrease in Th2 cytokines. C57BL/6 mice were treated with anti CD20 mAb or Rat IgG2a isotype control at 100 μg in 200 μl PBS by i.v. injection via tail vein prior to infection. Mice were infected with approximately 150 *T. muris* eggs. Mice were autopsied at day 35 p.i.. (A) Diagram of experimental design. (B) Worm burdens were assessed blindly after autopsy. (C-E) Sera were analysed using ELISA for parasite specific antibodies. (F-J) MLN cells were re-stimulated with parasite E/S antigen for 48 hours and cytokines in supernatants were determined using cytokine bead array (CBA). Data shows mean ± SEM, pooled from 3 independent experiments (B&F-K), representative of 3 independent experiments (C-E), males, *p<0.05, **p<0.01 ***p<0.001, Student’s t-test.

It has previously been shown that IgG1 antibodies are associated with resistance to *T. muris* infection, whilst IgG2c antibodies are related to susceptibility (32). Therefore, the levels of parasite specific antibodies in the sera of anti-CD20 treated mice and isotype control treated mice were compared. Mice depleted of B cells using anti-CD20 mAb failed to secrete IgG1 and IgG2c parasite specific antibodies, in contrast to control treated mice (Fig. 2C&D). Further, the levels of *T. muris* specific IgM antibodies in the sera of anti-CD20 treated mice were significantly lower than in the sera of the isotype control-treated mice (Fig. 2E). These effects of B cell depletion on antigen-specific IgG and IgM antibodies are consistent with reports in other model systems (33).

In common with other nematode parasites, hosts resistant to *T. muris* mount Th-2 immune responses characterised by the production of IL-4, IL-5, IL-9 and IL-13; in contrast, mice susceptible to *T. muris* mount a Th-1 type response, dominated by the release of IFN-γ (32, 34, 35). B cell depleted mice on a C57BL/6 background produced significantly lower Th2 cytokines, including IL-5, IL-9, and IL-13 compared to mice treated with the isotype control (Fig. 2F-H). Interestingly, the production of IFN-γ and IL-17 was similar in both groups (Fig. 2I&J) suggesting that B cells do not affect Th1 development but rather boost Th2 responses. These data are in keeping with the previous study by Leon, *et al*., which revealed that B cell depletion impaired the development Th2 in mice against *H. polygyrus* infection (15).

### 2.3 B cell depletion reduced the T_FH_ population and altered DC subsets in the MLN

The importance of B cells for the formation of T_FH_ has been shown previously in both mouse (15) and man (36). In keeping with these studies, the MLN T_FH_ population of anti-CD20 treated mice in C57BL/6 genetic background was significantly reduced compared to isotype control treated mice (Fig. 3A-C). Gating strategy was shown in Supplementary Fig. 1. In contrast, T_FH_ development in the spleen, distal to the site of infection, was not significantly affected by B cell depletion (Supplementary Fig.2), although the lack of statistical significance may be due to the high variability in the control mice.

**Fig. 3.**
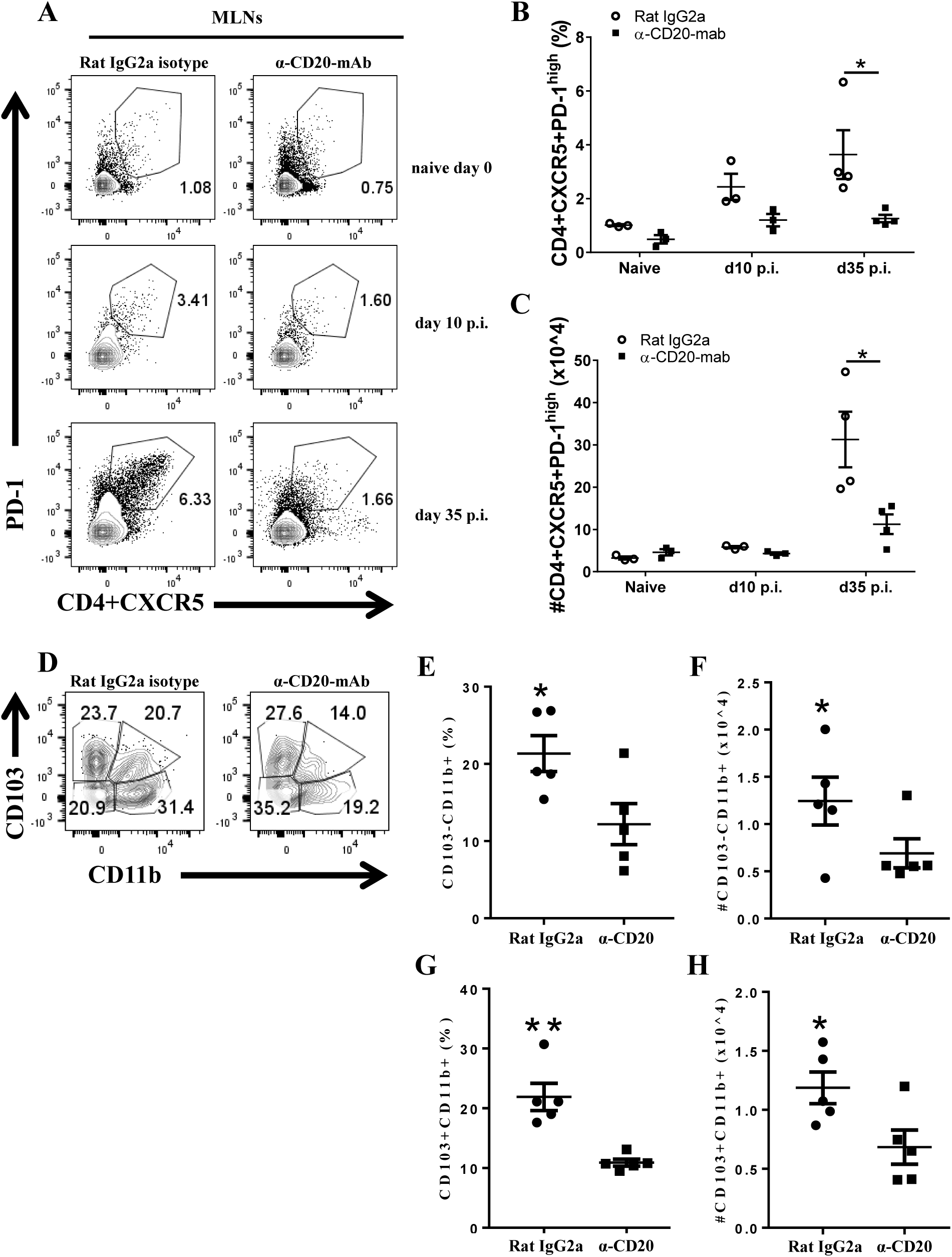
B cell depletion alters the T_FH_ population and DC subsets in MLNs. C57BL/6 mice were treated with anti CD20 mAb or Rat IgG2a isotype control at 100 μg in 200 μl PBS by i.v. injection via the tail vein. Mice were infected with approximately 150 *T. muris* eggs at day 7 post injection. Mice were re-injected with anti CD20 mAb or isotype control at 100 μg in 200 μl PBS by i.v. injection via the tail vein at day 10 p.i.. Mice were autopsied on day 0, day 10 and day 35 p.i. for T_FH_ and on day 35 p.i. for DC subsets. (A) Gating on CD4+CXCR5+PD-1^high^ to define T cells in MLNs. (B&C) Relative % and total cell number of T_FH_ in MLNs, respectively. (D-H) DC subsets in MLNs. (E&F) Relative percentage and total cell number of CD103-CD11b+ DCs, respectively. (G&H) Relative percentage and total cell number of CD103+CD11b+ DCs, respectively. Data shows mean ± SEM, from 1 experiment, n=3-5, males, *p<0.05, **p<0.01, Mann Whitney test.

Previous studies have suggested that CD11b^+^ DCs in MLNs might be important for promoting Th2 immune responses against *T. muris* infection (37). Because Th2 cytokines were significantly reduced in the MLNs from anti-CD20 treated mice during *T. muris* infection, DC subsets in MLNs were analysed (Fig. 3D-H). The gating strategy is shown in Supplementary Fig. 3. Total cell number of CD103^−^CD11b^+^ DCs (Fig. 3F) and CD103^+^ CD11b^+^ DCs (Fig. 3H) were significantly reduced in MLNs of anti-CD20 treated mice compared to isotype control treated mice.

### 2.4 B cell depletion from day 14 post infection also altered the worm burden

As depletion of B cells throughout the course of infection of C57BL/6 mice lead to susceptibility to *T. muris*, we wondered whether B cell depletion from day 14 p.i. would also alter the phenotype. Previous studies on the expulsion kinetics of *T. muris* have shown that the onset of worm expulsion occurs after day 12 p.i., with T cell activation occurring after the first 7 days (38). Therefore, B cells were depleted 2 weeks post *T. muris* infection to see if this still impacted on the ability to expel the parasite. The experimental design is shown in Fig. 4A. Surprisingly, the depletion of B cells from day 14 p.i. also impaired worm expulsion with significantly more parasites present at day 35 p.i. than seen in control-treated mice (Fig. 4B). T_FH_ cells and Th2 type cytokines, including IL-5, IL-9, and IL-13 were also significantly reduced in the absence of B cells (Fig, 4C-H). Interestingly, TNF-α was significantly increased in B cell depleted mice (Fig. 4I), whilst IFN-γ and IL-17A remained the same between groups (Fig. 4J&K). These data suggest that the B cell plays an important role in maintaining Th2 responses.

**Fig. 4.**
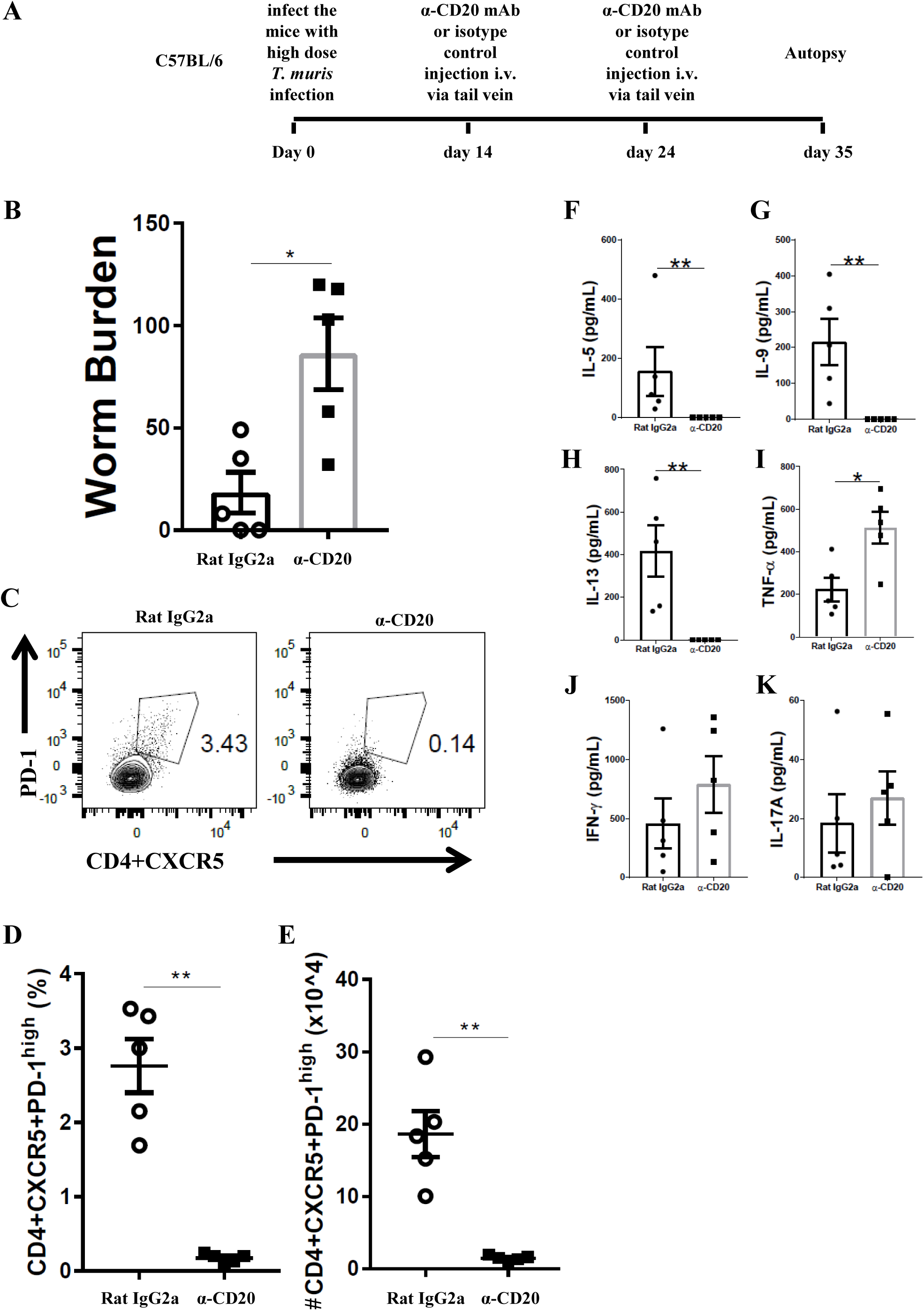
B cell depletion from day 14 p.i. also leads to susceptibility to *T. muris* infection. C57BL/6 were treated with anti CD20 mAb or Rat IgG2a isotype control at 100 μg in 200 μl PBS by i.v. injection via the tail vein 2 weeks post infection (p.i.). Mice were re-injected with anti CD20 mAb or isotype control at 100 μg in 200 μl PBS by i.v. injection via the tail vein on day 24 p.i. Mice were autopsied on day 35 p.i. (A) Diagram of the experiment design for B cells depletion 2 weeks post-infection. (B) Worm burdens were assessed blindly after autopsy. (C) Gating on CD4+CXCR5+PD-1^high^ to define T cells in MLNs. (D&E) Relative % and total cell number of T_FH_ in MLNs, respectively. (F-K) MLN cells were re-stimulated with parasite E/S antigen for 48 hours and cytokines in supernatants were determined using cytokine bead array (CBA). Data show mean ± SEM from 1 experiment, n=5, males, *p<0.05, **p<0.01, Mann Whitney test.

### 2.5 Anti-IFN-γ treatment partially restored IL-13 production in B cell depleted mice and rescued worm expulsion

In order to investigate whether the impaired resistance to infection in the absence of B cells was due to the reduced Th2 immune response or loss of parasite specific antibodies, B cell depleted mice were injected with anti-IFN-γ. Anti-IFN-γ treatment is a common strategy to promote resistance in susceptible mice (39). The experimental design for anti-IFN-γ injection plus B cell depletion is shown in Fig. 5A. As shown in Fig. 5B, anti-IFN-γ treatment restored resistance to *T. muris* in B cell depleted mice.

**Fig. 5.**
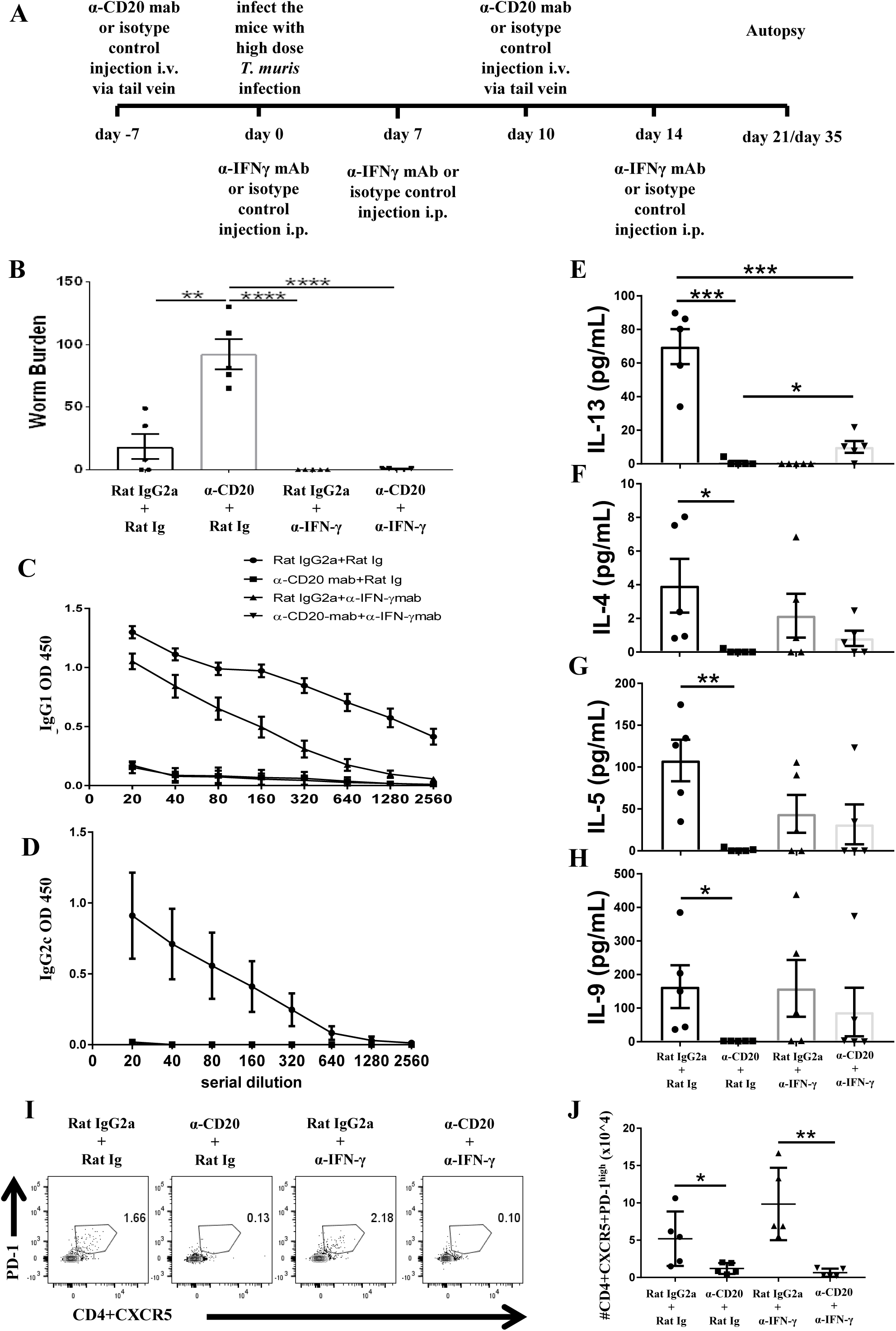
Anti-IFN-γ treatment restored resistance to *T. muris* infection and partially rescued the IL-13 response in B cell depleted mice in the absence of *T. muris* specific IgG1 antibodies and without preserving T_FH_ population. C57BL/6 mice were treated with anti CD20 mAb or isotype control at 100 μg in 200 μl PBS by i.v. injection via the tail vein. Mice were infected with approximately 150 *T. muris* eggs at day 7 post injection. Mice were re-injected with anti CD20 mAb or isotype control at 100 μg in 200 μl PBS i.v. injection via tail vein on day 10 p.i.. 1 mg of α-IFN-γ antibody or Rat Ig (as a control) was given on day 0, day 7, and day 14 p.i.. Mice were autopsied on day 21 p.i. and day 35 p.i.. (A) Diagram of the experimental design. (B) Worm burdens were assessed blindly after autopsy at day 35 p.i.. (C&D) *T. muris* specific IgG1 and IgG2c antibodies in the sera by day 21 p.i., respectively. (E-H) Cytokine analysis of re-stimulated MLN cells day 21 p.i.. (I) Gating on CD4+CXCR5+PD-1^high^ to define T_FH_ cells in MLNs day 21 p.i.. (J) Total numbers of T_FH_ cells in MLNs day 21 p.i.. Data shows mean ± SEM, from 1 experiment, n=5, males, *p<0.05, **p<0.01, ***p<0.001, ****p<0.0001, Anova.

As expected, blocking IFN-γ in the absence of B cells did not rescue the *T. muris* specific IgG1 response (Fig. 5C). T. muris specific IgG2c antibodies were not detected in the sera of anti-IFN-γ treated mice with intact B cells (Fig. 5D), consistent with a highly polarised Th2 immune response. In contrast, the isotype control mice produced both IgG1 and IgG2c antibodies (Fig. 5C&D). Consistent with our previous data, mice treated with anti CD20 mAb produced significantly lower Th2 cytokines, such as, IL-13, IL-4, IL-5 and IL-9 compared to isotype control treated mice (Fig. 5D-G) with the Th1 response unaffected (data not shown). Importantly, blocking IFN-γ significantly increased IL-13 production in mice depleted of B cells (Fig. 5D), although levels were still significantly lower than in isotype control treated mice (Fig. 5D). Although no significant difference was noted in the production of other Th2 type cytokines, there were trends towards increased *T. muris* specific IL-4, IL-5, and IL-9 after anti-IFN-γ treatment in B cell depleted mice compared to mice depleted of B cells but not IFN-γ (Fig. 5F-H). Taken together, these results show that in the absence of IFN-γ, B cells and IgG1 parasite specific antibodies are not important for *T. muris* expulsion on a C57BL/6 genetic background; thus without any competing IFN-γ partial restoration of the Th2 immune response is sufficient to render animals resistant to infection

Consistent with data in Fig 4, B cell depletion reduced the T_FH_ population in MLNs (Fig. 5I&J). T_FH_ cells have been associated with Th2 immunity due to their ability to secrete IL-4, a Th2 signature cytokine (33, 40, 41). Given that IL-13 production persisted in B cell depleted mice after anti-IFN-γ injection, we wondered whether the T_FH_ population was also altered. Anti-IFN-γ treatment did not restore the T_FH_ population in MLNs of anti-CD20 treated mice at day 21 p.i. (Fig. 5I&J), however it remains a possibility that the few TfH remaining were sufficient to induce the partial Th2 response.

### 2.6 B cells are essential in supporting Th2 immune responses against *T. muris* only in an IFN γ-rich environment

In the current study we found that when C57BL/6 mice were depleted of IFN-γ, the immune response became more highly polarised towards Th2 mice, evidenced by parasite specific IgG1 antibodies, in the absence of IgG2c. In this more polarised Th2 environment the depletion of B cells did not prevent worm expulsion. These data suggest that the important role played by B cells in supporting Th2 immune responses against *T. muris* is only necessary in IFN-γ-rich environments. Therefore, we decided to deplete B cells in BALB/c mice, which are naturally very resistant to *T. muris* and mount highly polarised Th2 immune responses in the absence of IgG2a/c and IFN-γ (31). The experimental design is shown in Fig. 6A. CD19+ cells in MLNs, spleen, and PerC of BALB/c mice were not detected after anti-CD20 mAb injection (Fig. 6B). In complete contrast to B cell depleted C57BL/6 mice, BALB/c mice were still able to expel the parasite in the absence of B cells (Fig. 6C). IgG1 and IgG2c parasite specific antibodies were undetectable in B cell depleted mice (Fig. 6D&E), and levels of *T. muris* specific IgM were significantly lower than in the sera of isotype control treated mice (Fig. 6F). As shown previously (31, 39), isotype control treated mice on BALB/c genetic background mounted a strong parasite specific IgG1 response but did not secrete *T. muris*-specific IgG2a/c (Fig. 3K), indicative of a highly polarised Th2 immune response.

**Fig. 6.**
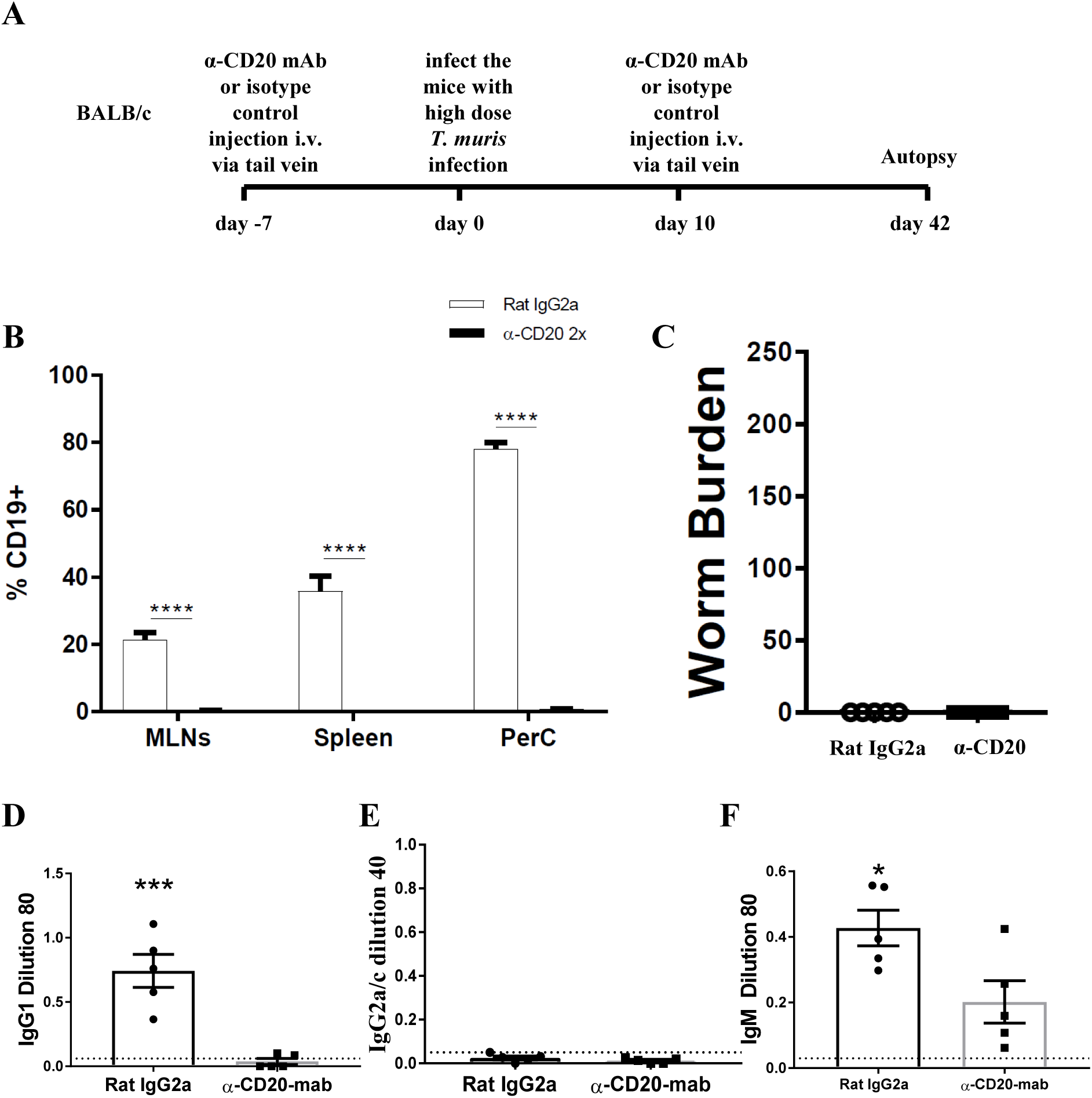
BALB/c mice were able to expel *T. muris* in the absence of B cells. BALB/c mice were treated with anti CD20 mAb or Rat IgG2a isotype control at 100 μg in 200 μl PBS by i.v. injection via tail vein prior to infection. Mice were infected with approximately 150 *T. muris* eggs and were autopsied at day 42 p.i.. (A) Diagram of experimental design. (B) CD19+ cells were assessed in MLNs, spleen, and peritoneal cavity (PerC) using flow cytometry. (C) Worm burdens were assessed blindly after autopsy. (D-F) Sera were analysed using ELISA for parasite specific antibodies. Data shows mean ± SEM, from 1 experiment, n=5, males. *p<0.05, ***p<0.001 ****p<0.0001, Student’s t-test.

Collectively, our data reveals that the important role played by the B cell in promoting resistance to *T. muris* infection in mixed Th1/Th2 cytokine settings is in supporting the development and maintenance of the Th2 immune response and is not related to antibody production. Importantly, the essential role played by the B cell in the protective immune response varied with genetic background and the degree of Th cell polarisation of the host. Thus, if IFN-γ is depleted from mixed Th1/Th2 settings, or the Th2 immune response is dominant, the B cell becomes redundant in the protective immune response.

## Discussion

*T. muris* infection drives different Th cell responses in different strains of mice (34). BALB/c, BALB/k, and NIH mice are very resistant to *T. muris* infection, expelling the worms by around day 18 post infection, while AKR mice are susceptible to infection, unable to expel the parasite and harbour patent chronic infections (42). C57BL6 and C57BL10 mice are also resistant to infection, but they expel the worms more slowly, between day 18 and 35 p.i. (43). Susceptible hosts mount predominantly Th-1 immune responses associated with the presence of IFN-γ and IL-12, whilst very resistant host strains mount strong Th-2 type polarised immune response characterised by cytokines such as IL-4, IL-5, IL-9 and IL-13, and very low levels of IFN-γ. The quality of the Th cell response in C57BL6 mice is less clearly polarised with the slower expulsion kinetic associated with a mixed Th1/Th2 phenotype and presence of IgG1 and IgG2c. Despite a good understanding of the Th cell response during *T. muris* infection, the key cellular contributions which support T helper polarization are still not well understood, nor how genetic background impinges on this.

This study aimed to investigate the role of B cells in immunity to *T. muris* during primary infection by using anti-CD20 mAb to deplete B cells. In order to understand whether the contribution of the B cell varied with the nature of the protective immune response, we used mice of two different genetic backgrounds: C57BL/6 and BALB/c. CD20 is specifically expressed on the surface of B cells from the pre-B cells stage to immature B cells, but then disappear when B cells differentiate to plasma cells (28). Adding antibody against CD20 inhibits the progression of B cells from the G1 phase into S/G2+ M stages (44), resulting in the inhibition of B cell differentiation, antibody production, and inducing B cell apoptosis (45). Using anti-CD20 mediated depletion of B cells, antigen specific class switched antibodies are ablated and IgM antibodies are significantly reduced (41, 46). These finding are in keeping with the finding of current study and enable an assessment of the importance of parasite-specific IgG1 antibodies in the expulsion of *T. muris*.

Our data reveal that antibody is not essential for the expulsion of *T. muris*, with both B cell depleted BALB/c mice and B cell depleted C57BL/6 mice also depleted of IFN-γ, able to clear infection. These data also exemplify that the mechanism of immunity to *T. muris* in BALB/c mice is entirely B cell independent. However, our data do identify an important and significant antibody-independent role for the B cell in promoting immunity to infection in mouse strains, such as C57BL/6, which are not highly polarised towards a Th2 immune response. Thus, the importance of the B cell in resistance to infection is dependent on genetic background, and represents a key consideration when interpreting data from other experimental models. Further, given that protective immunity was lost when B cells were depleted both throughout the time course of infection and only after the first two weeks of infection, our data suggest that the role played by the B cell is in supporting both the development and maintenance of Th2 immune responses.

Mechanistically, B cell depletion correlated with a significant reduction in CD4+CXCR5+PD-1^high^ cells. CD4+CXCR5+PD-1^high^ cells are recognised as T_FH_ and are known to be able to produce the Th2-type signature cytokine; IL-4 (47). Thus, these data are consistent with previous studies showing that B cells are important in regulating Th2 immune responses (10, 14, 15, 48–51) and maintaining T_FH_ development (15, 52). Interestingly, T_FH_ populations in both MLN and spleen of BALB/c mice were not affected after B cell depletion. Further, the presence of T_FH_ has been associated with Th2-type immune responses in *H. polygyrus* infection (15). Although in the current study, depletion of B cells in mice also treated with anti IFN-γ restored expulsion of *T. muris* without rescuing the T_FH_ response, the few T_FH_ cells remaining may be sufficient to support the partially restored Th2 response. Thus it remains possible that T_FH_ cells are essential in building strong Th2 responses in the absence of IFN-γ.

The importance of B cells in immunity to *T. muris* has previously been proposed using µMT mice (22) on a C57BL/6 background which are susceptible to infection unless Th1 responses are inhibited using anti-IL-12. These data are keeping with the current study supporting a role for B cells as accessory cells promoting and maintaining Th2 responses rather than antibody producers. Antibody independent expulsion of *T. muris* is also evidenced by the fact that primed CD4+ T cell transfer to SCID mice is sufficient to support worm expulsion in the absence of antibody (18).

More broadly, the role of B cells in immunity to gastro intestinal nematodes in general has been debated at length (53–56). Proposed mechanisms include production of antibody (57), promoting and maintaining of primary and memory Th2 cells (10, 14) via their ability to produce cytokines, especially IL-4 (11, 53, 58), and/or by expressing co-stimulatory molecules, including OX40L and CD40 (13, 59). A variety of experimental approaches have been used including the use of transgenic B cell deficient mice (10, 22, 57, 60), FcγR deficiency (61, 62), passive immunization (22, 63, 64) and maternal antibody transfer (54, 65); and a variety of conclusions have been drawn. IgG1 antibodies were shown to be essential for worm expulsion against *H. polygyrus* infection based on previous studies using AID mice, which retain a secretory IgM response, on C57BL/6 genetic background (57). Antibody was shown to be essential for worm expulsion against *H. polygyrus* infection via antibody-dependent cell-mediated cytotoxic (ADCC) mechanisms (66). However, ADCC mechanisms do not play a critical role in immunity to *T. muris* infection as FcγR deficiency mice are still able to expel the parasite (62).

More recently, two studies have strongly suggested a role for B cells in directing T cell polarisation towards Th2 at the time of antigen presentation (15, 67). Leon *et al*., showed that B cell depletion in *H. polygyrus*-infected C57BL/6 mice prevented the migration of DCs into B cell areas and impaired the Th2 immune response (15). Lymphotoxin, produced by B cells, is important in the control of CXCL13 expression (15), a chemokine expressed by follicular dendritic cells (68) and marginal reticular cells in the peri and interfollicular regions between B cell follicles (69) and that is essential to attract CXCR5-expressing cells towards the B cell area (15). Thus by treating C57BL/6 mice with anti-CXCL13, Leon *et al.*, (15) also revealed that CD4 T cells and DCs accumulated in the T cell area rather than in perifollicular region and that IL-4+ Th2 cells in mesenteric lymph nodes (MLNs) were significantly reduced. In support of a mechanism involving the movement of DCs and CD4 T cells to the B cell area to allow CD4 T cells to be educated by B cells towards a Th2, recent studies have revealed that transgenic mice in which CXCR5 depletion was specifically restricted to DCs become susceptible to *T. muris* with very few DCs detected in the B cell area compared to control mice (67).

In addition to the proposed role for B cells in educating the DC-CD4 T cell conversation suggested by Leon *et al.*, (15) and Bradford *et al.*, (67), the importance of CD11b+ DCs in MLNs for Th2 polarisation has been shown in previous studies, against both *T. muris* (37) and *S. mansoni* (70) infection. In this context, in the current study we observed reduced numbers of CD11b+ DCs in B cell depleted C57BL/6 mice and this may contribute to the reduced Th2 immune response and increased susceptibility of anti-CD20 treated mice to *T. muris* infection. Although previous studies have indicated the importance of B cells in regulating the development of DC subsets in secondary lymphoid organs (50, 71, 72), further studies are required to investigate how the absence of B cells reduced the number of CD11b+ DCs in the MLN.

Overall, we present a study showing that, irrespective of host genetic background, antibody is not essential in the expulsion of *T. muris*. However, our data do identify an essential role played by the B cell on certain genetic backgrounds where IFN-γ rich, mixed Th1/Th2 cell responses are observed, and strongly suggest that the B cell is able to adopt multiple functions in inducing and maintaining protective Th2 responses during gastro intestinal nematode infection.

## Material and Methods

### 1. Animals

C57BL/6 and BALB/c mice were purchased from Envigo, UK and were maintained in ventilated cages in the Biological Services Facilities (BSF) of the University of Manchester. Mice were housed in the facility at least 7 days prior to experimentation and were infected at 6-8 weeks old with *T. muris* by oral gavage.

### 2. B cells depletion and infection

To assess the importance of B cells for worm expulsion in different strains, C57BL/6 and BALB/c mice were split into 2 groups of 4-5 mice: anti-CD20 mAb and isotype control treated mice. C57BL/6 or BALB/c mice were treated with 100μg in 200μl PBS of anti-CD20 mAb (5D2, Genentech) or isotype control (Rat IgG2a, Biolegend) i.v. injection via tail vein. Mice were infected with approximately 200 T. muris eggs at day 7 post injection. Mice were re-injected with anti-CD20 mab or isotype control at day 10 p.i. C57BL/6 mice were autopsied at day 35 p.i., whilst BALB/c mice were autopsied at day 42 p.i. For anti-IFN-γ, C57BL/6 mice were split into 4 groups of 5 mice: isotype control of anti-CD20 mAb + isotype control of anti-IFN-γ mAb (rat Ig), anti-CD20 mAb + isotype control of anti-IFN-γ mAb, isotype control of anti-CD20 mAb + anti-IFN-γ mAb, anti-CD20 mAb + anti-IFN-γ mAb. C57BL/6 mice were treated with 100μg in 200μl PBS of anti-CD20 mAb (5D2, Genentech) or isotype control (Rat IgG2a, Biolegend) i.v. injection via tail vein. Mice were infected by oral gavage with approximately 200 *T. muris* eggs at day 7 post injection. Mice were re-injected with anti-CD20 mab or isotype control at day 10 p.i. 1 mg of α-IFN-γ antibody or rat IgG1 (as a control) antibody was given at day 0, day 7, and day 14 p.i. Mice were euthanised by CO2 followed by autopsy at day 21 or day 35 p.i..

### 3. *Trichuris muris* maintenance and the preparation of E/S proteins

All protocols to maintain the parasite and to prepare the E/S were as previously described (73). Briefly, the parasite was passaged through SCID mice that are susceptible to *T. muris* infection. SCIDs received a high dose of approximately 300 *T.muris* embryonated eggs and at approximately day 35 p.i. the large intestine was collected to produce adult E/S. Guts were collected and then longitudinally split open before washed in warmed 5x penicillin/streptomycin in complete RMPI 1640 medium (500U/ml penicillin, 500μg/ml streptomycin, 500ml RMPI 1640 medium). Adult worms were then carefully pulled from the gut using fine forceps and transferred to a 6 well plate containing 4ml warmed 5x pen/strep in RMPI 1640 medium. Plates were incubated in a moist humidity box for 4 hours at 37°C to collect 4 hr E/S. For overnight E/S, adult worms were then split into 2 wells containing fresh medium and incubated again in a humidity box at 37°C overnight. Supernatant from 4 hour and 24 hour incubations was collected and centrifuged at 2000g for 15 minutes. *T. muris* eggs from adult worms were resuspended in 40ml deionised water and filtered through a 100μm nylon sieve before transferring to a cell culture flask. To allow embryonation, eggs were stored in darkness for approximately 8 weeks and then stored at 4°C. SCID mice were subsequently infected with a high dose infection to determine infectivity of each new batch of eggs. Thus, larvae were counted at around d14 p.i. and the number of larvae counted expressed as a % over the number of eggs given to determine the infectivity of the egg batch. All E/S supernatant was filter sterilised through a 0.2μm syringe filter (Merk). E/S was concentrated using an Amicon Ultra-15 centrifugal filter unit (Millipore) by spinning at 3000g for 15 minutes at 4°C. E/S was dialysed against PBS using Slide-A-Lyzer Dialysis Cassettes, 3.500 MWCO (Thermo Science) at 4°C. The concentration of E/S was measured using the Nanodrop 1000 spectrophotometer (Thermo Fisher Science) and aliquoted before storing at −80°C.

For high dose *Trichuris muris* infection, approximately 3-4 ml of egg suspension was transferred to a universal tube and topped up with deionised water before centrifuging for 15 minutes at 2000g. Pelleted eggs were washed with deionised water, resuspended and only embryonated eggs were counted. Eggs were concentrated or diluted with deionised water, depending on the egg count. For example, eggs with 100% infectivity, approximately 50 eggs per 50μl were counted. Mice were then infected with 200μl by oral gavage to infect mice with approximately 200 eggs.

### 4. Worm burden of *Trichuris muris*

During autopsy, the caecum and proximal colon were collected and stored at −20°C before analysis. Before worm count, the intestine was thawed at room temperature and cut longitudinally using blunt ended scissors and the epithelium was scraped using curved forceps in a petri dish. Worms were counted blindly under a dissecting microscope (Leica).

### 5. Preparation of single cell suspensions for fluorescence activated cell sorting (FACS)

Mesenteric lymph nodes (MLNs), spleen, blood and bone marrow were collected and prepared for FACS staining. Lymph nodes and spleen were squeezed through a 70μm nylon cell strainer (Fisher Scientific) and cells were pelleted by centrifugation at 1500 rpm for 5 minutes. The supernatant was removed and the pelleted cells were resuspended in 500μl to 1 ml of Red Blood Cell Lysing Buffer Hybri-Max™ (Sigma-Aldrich) for 30 seconds to 1 minute before adding 10 ml 1xPBS. Cells were pelleted by centrifugation at 1500 rpm for 5 minutes and resuspended in 1 ml of complete RPMI 1640 medium. Cells were counted on a CASY cell counter (Scharfe System).

Approximately 50μl of blood were placed into 1.5 ml eppendorf tube containing 50μl 0.5M EDTA and stored on ice before analysis. 500 μl of Red Blood Cell Lysing Buffer Hybri-Max™ (Sigma-Aldrich) were added and samples were incubated for 5 minutes at room temperature. 1 ml of 1xPBS were added and cells were pelleted by centrifugation at 1500 rpm for 5 minutes. Red blood cell lysis process was repeated twice before cells were resuspended in 1 ml of complete RPMI 1640 medium.

Femurs and tibias were collected at autopsy and after removing any remaining tissue without damaging the bone integrity, the bone was placed on ice until ready to process. Bone was transferred to 70% ethanol for 2-3 minutes and then rinse in 3 changes of 1xPBS. In a petri dish, both ends of bone were cut and the bone was flushed gently with 1xPBS using a 3cc syringe and a 23 ga needle. To break up the clumps, the marrow was sucked up and was gently pushed back. Single cells suspension was filtered through 100μm nylon cell strainer (Fisher Scientific) and cells were pelleted by centrifugation at 1500 rpm for 5 minutes and resuspended in 1 ml of 1xPBS. The supernatant was removed and the pelleted cells were resuspended in 500μl of Red Blood Cell Lysing Buffer Hybri-Max™ (Sigma-Aldrich) for 1 minute before adding 10 ml 1xPBS. Cells were pelleted by centrifugation at 1500 rpm for 5 minutes and resuspended in 1 ml of complete RPMI 1640 medium.

### 6. Cell surface markers

Cells from MLNs, spleen, blood and bone marrow were stained for live dead (Zombie UV, Biolegend) and Fc block (eBiosciences) prior to cell surface cellular markers staining. Samples were read on a BD LSR Fortessa flow cytometer (BD Biosciences) and data was analysed using FlowJo X (Tree Star, Inc).

Cell surface markers: anti-B220 (RA3-6B2); anti-CD19 (6D5); anti-CD3ε (17A2); anti-CD4 (RM4.5); anti-CD8α (53-6.7); anti-CD279 (PD-1) (29F.1A12); anti CD185/CXCR5 (L138D7); anti-CD93 (AA4.1); anti-CD25 (3C7); and anti-CD117/c-kit (2B8) were purchased from Biolegend. Anti-CD43 (S11); anti-CD103 (2E7); anti-CD11c (N418); anti-CD317/PDCA-1 (927); anti CD11b (M1/70); anti-CD64 (X54-5/7.1); anti-I-A/I-E (M5/114.15.2); and anti-CD23 (B3B4) were purchased from BD Biosciences. Anti-CD45 (30-F11); anti-ly6G (RB6-8C5); anti-NK.1 (PK136); anti-Ter119 (Ter-119) purchased from eBiosciences.

### 7. Quantification of parasite specific IgG1, IgG2a/c and IgM

To detect IgG1, IgG2a/c and IgM specific *T. muris*, an enzyme linked immunosorbant assay (ELISA) was completed. Blood was collected from mice at autopsy and serum was isolated by centrifuging samples for 10 minutes at 15,000g at room temperature. 96 well immunoGrade plates (BrandTech Scientific, Inc) were coated with 5μg/ml *T. muris* diluted in 0.05M carbonate/bicarbonate buffer and incubated on plates overnight at 4°C. Plates were washed using a Skatron Scan Washer 500 (Molecular Devices, Norway) 3 times with 0.05% Tween 20 (Sigma) in PBS (PBS-T). Non-specific binding were blocked with 100μl 3% bovine serum albumin (BSA) (Melford Laboratories)/ PBS at 37°C for 45 minutes. Plates were washed 3 times with PBS-T and 50μl of double diluted serum in PBS (1:20, 1:40, 1:80, 1:160, 1:320, 1:640, 1:1280, 1:2560) was added to plates and incubated for 60 minutes. Plates were then washed 3 times with PBS-T and 50μl of either biotinylated rat anti-mouse IgG1 (1:500, BD Bioscience), rat anti-mouse IgG2a (1:1000, BD Bioscience) or rat anti-mouse IgM (1:500, BD Bioscience) was added to wells and incubated for 60 minutes. C57BL/6 mice do not make IgG2a, but IgG2c. However, the rat anti-mouse IgG2a antibody also recognises mouse IgG2c, the IgG2a equivalent. Plates were then washed with PBS-T 3 times before the plates were incubated with 75μl Streptavidin peroxidase (1:1000, Sigma) for 60 minutes. Plates were washed 3 times with PBS-T before the colour were developed using 100μl 0.03% hydrogen peroxidase activated ABTS (10% 2,2’azino 3-thyl benzthiazoline in 0.045M citrate buffer). Plates were read at 450nm on a VersaMax Microplate reader (Molecular Devices).

### 8. Cytokine analysis

During autopsy, mesenteric lymph nodes were isolated and collected in complete RPMI 1640 medium. The tissue were squeezed through a 70μm nylon cell strainer (Fisher Scientific) and cells were pelleted by centrifugation at 1500 rpm for 5 minutes. The supernatant was removed and the pelleted cells were resuspended in 500μl to 1 ml of Red Blood Cell Lysing Buffer Hybri-Max™ (Sigma-Aldrich) for 30 seconds to 1 minute before adding 10 ml 1xPBS. Cells were pelleted by centrifugation at 1500 rpm for 5 minutes and resuspended in 1 ml of complete RPMI 1640 medium. Cells were counted on a CASY cell counter (Scharfe System) and then diluted to a concentration of 1×10^7^ cells/ml. 100μl of cells were plated out into a 96 well flat bottom plate. Cells were restimulated with 100µl 4 hr *T. muris* E/S at 100μg/ml (to give a final concentration of E/S in the cell culture of 50µg/ml). Cells were incubated for 48 hours at 37°C and 5% CO2. To collect the supernatant, plates were centrifuged at 1400g for 5 minutes and the supernatant was stored at −20°C.

The cytokines IL-4, IL-5, IL-6, IL-9, IL-10, IL-17, IL-13, TNF and IFN-γ were detected in supernatant by Cytometric bead assay (CBA). 12.5μl of supernatant from *T. muris* E/S stimulated cells was added to a 96-well round bottom plate. A capture bead cocktail (BD Bioscience), containing beads for each cytokine, was diluted in capture diluent (BD Bioscience) and 12.5μl was added to each well before incubating on a rocker at room temperature for 1 hour. 12.5μl of detection beads (BD Bioscience), diluted in detection reagent (BD Bioscience) was added to each well and incubated again on a rocker at room temperature for 1 hour. Plates were washed and resuspended in 70μl wash buffer (BD Bioscience). Cytokines were measured on a MACSQuant Analyser (Miltenyi Biotec) and analysed using the FCAP array software in reference to a standard curve.

### 9. Statistical analysis

Statistical analysis was performed using Prism4 (Graph-Pad software Inc., La Jolla, CA). The significant differences between two groups (P<0.05) were analysed with the t-test or Mann Whitney test, depend on the n-size and the distribution of samples. For multiple groups, the significant differences were analysed by Anova.

### 10. Ethics statement

All experiments were approved by The University of Manchester Local Ethical Review Committee and were performed in accordance with the UK Home Office Animals (Scientific Procedures) Act 1986, under the home Office project licence number 70/8127

## Acknowledgment

This study was funded by the Indonesian endowment fund for education. We would like to thank Rebecca Dookie for helping us with i.v. injection, Genentech (San Francisco, CA) for their gift of purified anti-CD20 mAb (clone-5D2), and Gareth Howell from Flow cytometry facility at the University of Manchester.

---

**Supplementary Fig.1.**
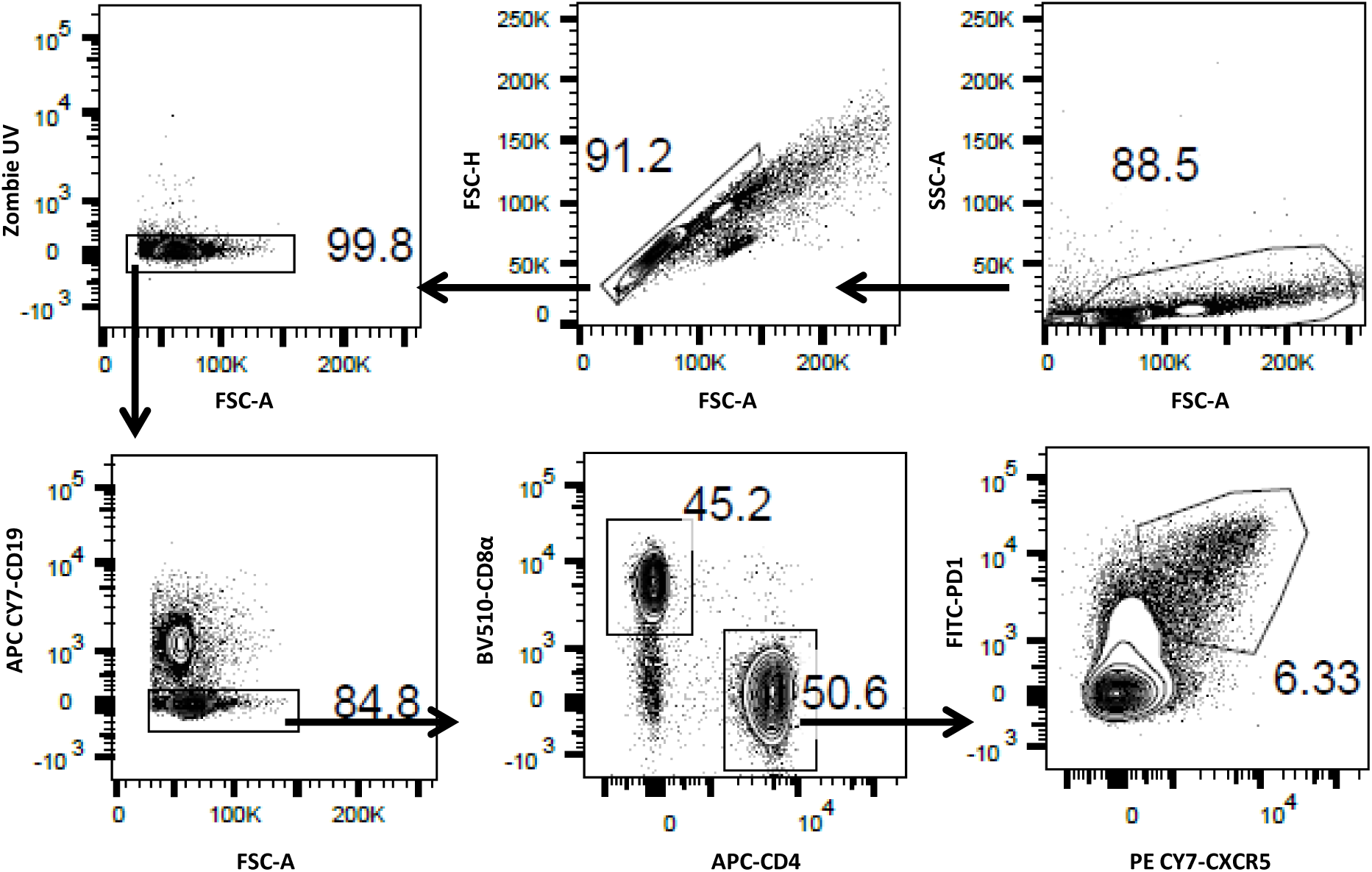
Flow cytometry gating strategy for T_FH_. T_FH_ cells are defined as CD4+CXCR5+PD-1^high^

**Supplementary Fig. 2.**
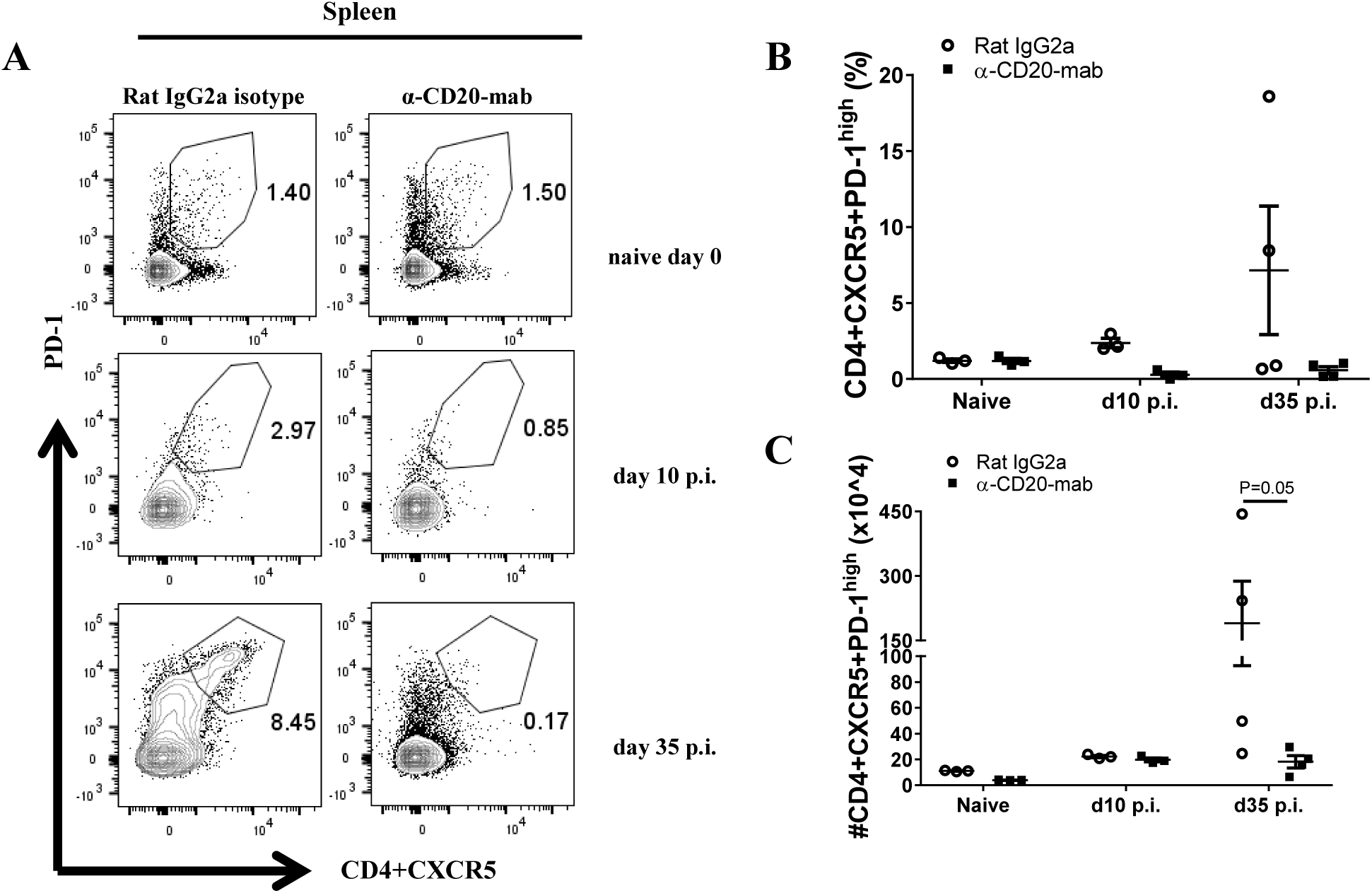
T_FH_ population in the spleen of anti-CD20 mAb treated mice. C57BL/6 mice were treated with anti CD20 mab or isotype control 100 μg in 200 μl PBS i.v. injection via tail vein. Mice were infected with approximately 150 *T. muris* eggs at day 7 post injection. Mice were re-injected with anti CD20 mab or isotype control 100 μg in 200 μl PBS i.v. injection via tail vein by day 10 p.i. Mice were autopsied by day 0, day 10 and day 35 p.i. Gating on CD4+CXCR5+PD-1^high^ to define T_FH_ cells in spleen (A). (B&C) Relative % and total cell number of T_FH_ in the spleen, respectively. Data show mean±SEM, from 1 experiment, males., *p<0.05, Mann whitney U test

**Supplementary Fig.3.**
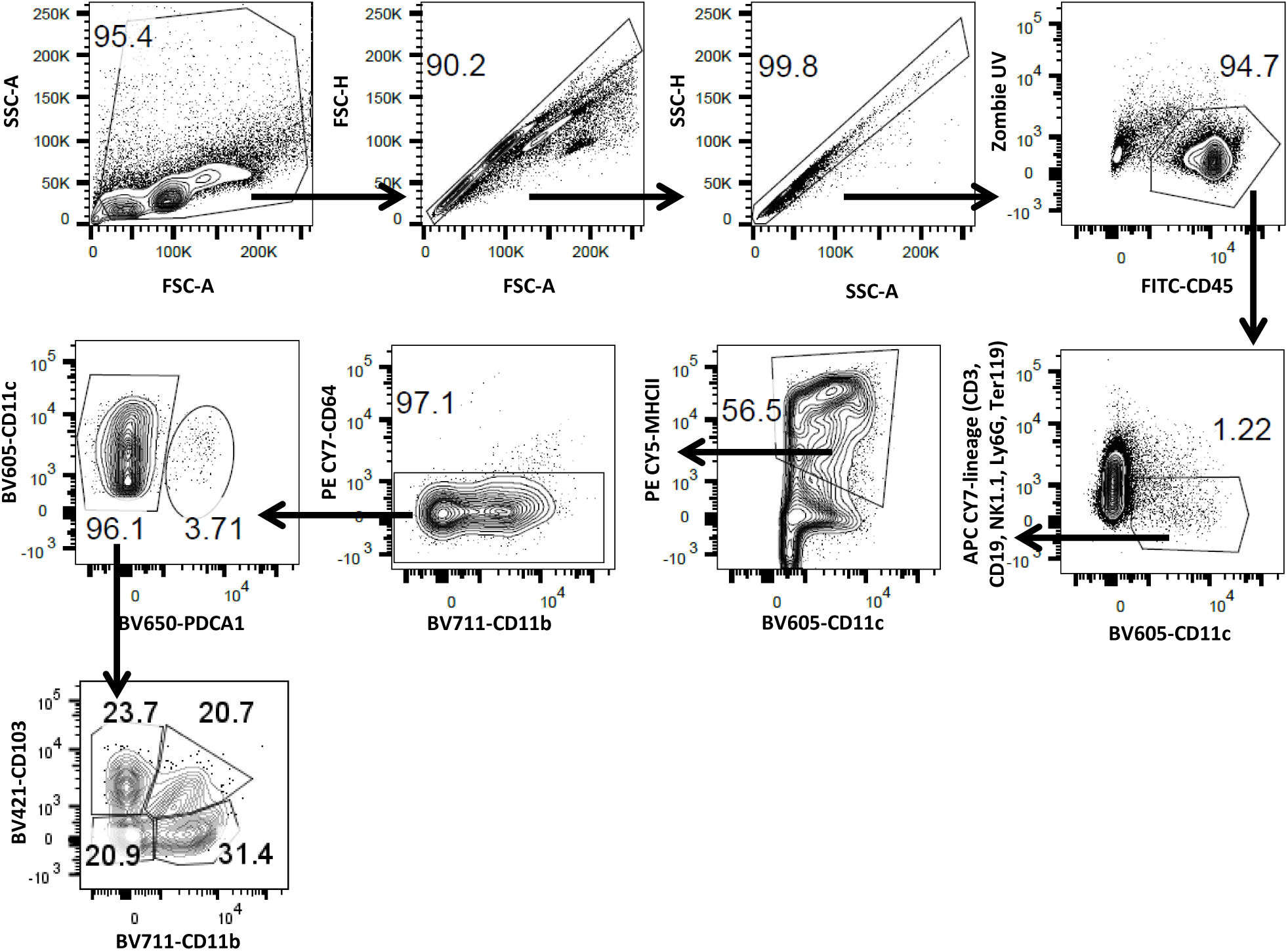
Flow cytometry gating strategy for DC subsets. DCs are defined as CD45+lineage-CD11c+MHCII+CD64-. Two main DCs populations: conventional DCs (cDCs) and plasmacytoid DCs (pDCs). Conventional DCs (CD45+lineage-CD11c+MHCII+CD64 PDCA1-) were divided into migratory DCs and resident DCs. Migratory DCs was consisted of 4 subpopulations: CD103+CD11b-, CD103+CD11b+, CD103-CD11b+ and CD103-CD11b-. Plasmacytoid DCs:CD45+lineage-CD11c+MHCII+CD64-PDCA1+

**Supplementary Fig. 4.**
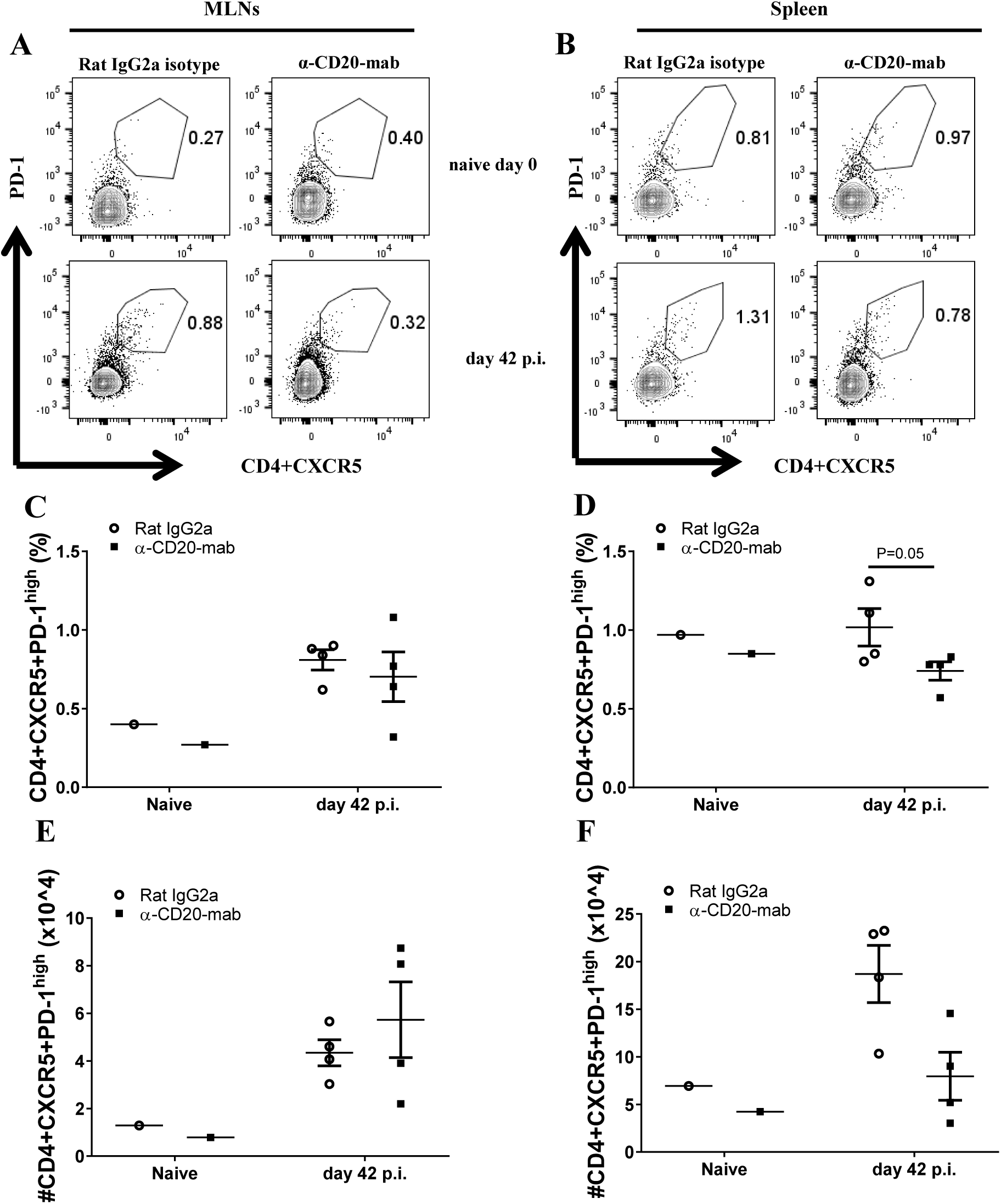
Tfh population in the MLNs and spleen of anti-CD20 mab treated mice on a Balb/c genetic background were not affected. Balb/c mice were treated with anti CD20 mab or isotype control 100 μg in 200 μl PBS i.v. injection via tail vein. Mice were infected with approximately 150 *T. muris* eggs at day 7 post injection. Mice were re-injected with anti CD20 mab or isotype control 100 μg in 200 μl PBS i.v. injection via tail vein by day 10 p.i. Mice were autopsied by day 0 and day 42 p.i. Gating on CD4+CXCR5+PD-1^high^ to define Tfh cells in MLNs (A) and spleen (B). (C&D) Relative % of Tfh in MLNs and spleen, respectively. (E&F) Total Tfh in MLNs and spleen, respectively. Data show mean±SEM, from 1 experiment, males.

